# The effector TepP mediates the recruitment and activation of Phosphoinositide 3 Kinase on early *Chlamydia trachomatis* vacuoles

**DOI:** 10.1101/132282

**Authors:** Victoria Carpenter, Yi-Shan Chen, Lee Dolat, Raphael H. Valdivia

## Abstract

*Chlamydia trachomatis* delivers multiple Type 3 secreted effector proteins to host epithelial cells to manipulate cytoskeletal functions, membrane dynamics and signaling pathways. TepP is the most abundant effector protein secreted early in infection but its molecular function is poorly understood. In this report, we provide evidence that TepP is important for bacterial replication in cervical epithelial cells, the activation of Type I IFN genes, and the recruitment of Class I phosphoinositide 3 kinases (PI3K) and the signaling adaptor protein CrkL to nascent pathogen-containing vacuoles (inclusions). We also show that TepP is a target of tyrosine phosphorylation by Src kinases but these modifications do not appear to influence the recruitment of PI3K or CrkL. The translocation of TepP correlated with an increase in the intracellular pools of phosphoinositide 3,4,5 triphosphate but not the activation of the pro-survival kinase Akt, suggesting that TepP-mediated activation of PI3K is spatially restricted to early inclusions. Furthermore, we linked PI3K activity to the dampening of transcription of Type I IFN induced genes early in infection. Overall, these findings indicate that TepP can modulate cell signaling and potentially membrane trafficking events by spatially restricted activation of PI3K.

## INTRODUCTION

*Chlamydia trachomatis* is an obligate intracellular bacterial pathogen of significant socio-economic and medical importance. *C. trachomatis* is the leading causative agent of preventative blindness worldwide and the most prevalent sexually transmitted infection (STI) in the western world (1, 2). *C. trachomatis* undergoes two main developmental transitions with an infectious form, the elementary body (EB), and a replicative form, the reticulate body (RB). Both RB and EB forms of the pathogen manipulate host cellular functions by delivering Type 3 Secretion (T3S) effector proteins directly into the target cells’ membranes and cytoplasm (3).

The *Chlamydia* T3S system shares many functional features with T3S systems from other Gram negative bacteria, including the requirement for accessory chaperones that stabilize effectors and enhance their secretion (3-5). One of these T3S chaperones, Slc1, interacts with and enhances the secretion of multiple EB effectors (6). For instance, the effector Tarp (Translocated actin recruiting phosphoprotein) is delivered into epithelial cells within 5 minutes of EB attachment and phosphorylated at tyrosine residues (7-9). Multiple proteins with Src homology 2 (SH2) domains can bind *in vitro* to peptides representing the phosphorylated forms of Tarp (10, 11). This includes the E3 ligase Cbl, the Rac1 exchange factor Vav2, the p85 regulatory subunit of phosphoinositide 3 kinase (PI3K), the signaling adaptors Shc1, Nck2 and CrkL, and the kinase Syk (12). Various tyrosine kinases can phosphorylate Tarp *in vitro* and *in vivo*, including Src, Abl and Syk (7, 8). Tarp nucleates F-actin assembly, independently of its tyrosine phosphorylation status, through its F-actin-binding domain and interaction with the Arp2/3 complex, and by activating focal adhesion kinases (FAK) through molecular mimicry of the cofactor paxillin (9, 11, 13-16). Not surprisingly, this multifunctional effector is important for *Chlamydia* infection as microinjection of anti-Tarp antibodies into epithelial cells or expression of dominant negative Tarp constructs in *Chlamydia* inhibit bacterial invasion (11, 17).

A second Slc1-dependent effector is the <u>T</u>ranslocated <u>e</u>arly <u>p</u>hospho-<u>P</u>rotein (TepP). On a molar basis, TepP is one of the most abundant *Chlamydia*-specific proteins found within EBs (18). Once translocated into the host cell TepP is phosphorylated by host cell kinases at multiple tyrosine and serine residues (6). Two tyrosine phosphorylation sites generate consensus pYxxP motifs that provide docking site for proteins with SH2 domains including two spliceforms of the signaling adaptor protein Crk (Crk I-II) (reviewed in (19)). Indeed, CrkI and CrkII co-immunoprecipitate with TepP and are recruited to nascent *C. trachomatis* inclusions in a TepP-dependent manner (6). A comparison of the transcriptional responses of epithelial cells to infection with a *tepP* deficient mutant and TepP over-expressing strains indicated a role for TepP in the induction of a subset of genes associated with Type I IFN responses, including interferon-induced peptides with tetratricopeptide repeats (*IFIT* (20)) (6).

To address the mechanism by which TepP modulates host cellular functions, we identified host proteins that associate with TepP during the early stages of bacterial invasion and establishment of inclusions. We determined that CrkL and Class I phosphoinositide 3 kinases (PI3K) (21) are the major proteins that co-purify with TepP and that these proteins are recruited to nascent inclusion in a TepP-dependent manner. Furthermore, TepP induces the activation of PI3K on internal membranes and nascent inclusions, to generate phosphoinositide 3,4,5 triphosphate (PIP3), without activating canonical PI3K signaling at the plasma membrane.

## RESULTS

### CrkL and PI3K co-purify with TepP translocated during infection

TepP is phosphorylated at multiple tyrosine residues upon delivery into host cells (6) and may directly recruit Src-homology 2 (SH2) and phosphotyrosine binding (PTB) domain-containing proteins to assemble novel host cell signaling complexes (22, 23). To identify *Chlamydia* and host proteins associated with TepP-containing signaling complexes, we infected A2EN endocervical epithelial cells with the *tepP* null mutant strain CTL2-M062G1, and variants transformed with either an empty plasmid (pVec) or a plasmid expressing TepP-FLAG (pTepP). After 4 hours, infected cells were lysed under non-denaturing conditions and subjected to immunoprecipitation (IP) with anti-FLAG antibodies. All proteins in the IP were digested with trypsin and the resulting peptides identified by liquid chromatography coupled to tandem mass spectrometry (LC-MS/MS). The major human proteins co-purifying exclusively with TepP-FLAG included the catalytic (p110α, p110β) and regulatory (p85α and p85β) subunits of PI3K, CrkL, and glycogen synthase kinase (GSK) (Fig. 1A and Table S1). The specificity of these interactions was verified by immunoblot analysis of subsequent IPs. CrkL, GSK and both PI3K subunits (p110 and p85) co-precipitated with TepP during infection (Fig. 1B). Reciprocal IP of CrkL and p110α co-precipitated phospho-TepP from infected cells (Fig. 1C and D), validating the specificity of these interactions.

**Figure 1:**
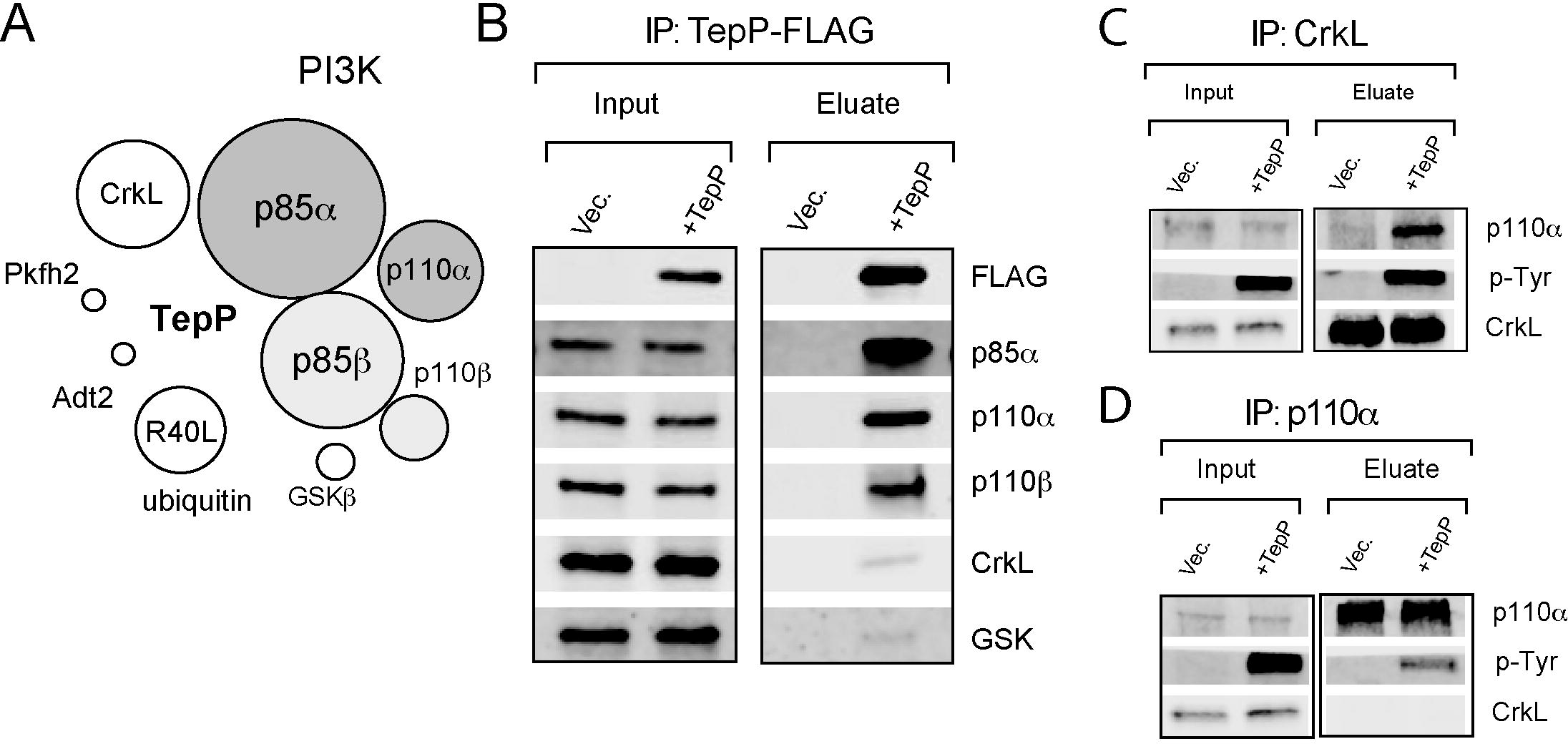
TepP forms complexes with PI3K and CrkL in infected cells. **(A)** Schematic representation of epithelial proteins that associate with TepP during infection. A2EN cells were infected with *C. trachomatis* expressing TepP-FLAG for 4h and cell lysates subjected to immunoprecipitation (IP) with anti-FLAG antibodies. Bound proteins were digested with trypsin and the resulting peptides identified by mass spectrometry (MS). Size of circles correspond to the relative number of peptides identified for each protein by LC-MS/MS. (**B)** Immunoblot validation of TepP-interacting proteins. A2EN cells infected for 4h with CTLM062G1 transformed with a TepP-FLAG expression plasmid or vector only control and cell lysates subjected to IP with anti-FLAG antibodies. Bound proteins were detected by immunoblotting with specific antibodies. **(C-D)** Reciprocal co-IP of CrkL (C), PI3K (D) and TepP. A2EN cells were infected with CTLM062G1 strains as in (B) and 4 h post infection CrkL and p110α were IP. Co-IP of oproteins was assessed by immunoblot analysis.

### TepP-deficient mutants fail to recruit PI3K and CrkL to early inclusions

To perform a clean phenotypic analysis of TepP mutants, we disrupted the *tepP* gene in CTL2 with a gene encoding β lactam resistance (*bla*) by Group II intron mediated gene insertion (TargeTron^TM^) (24). TepP could not be detected in insertional mutants by immunoblot analysis (Fig. 2A). The resulting *tepP::bla* (Δ*tepP*) strain behaved as the CTL2-M062G1 *tepP* nonsense mutant in that it failed to induce the expression of Interferon induced peptides with tetratricopeptide repeats (*IFIT*) genes (Fig. 2B) that are typically activated during the early stages of infection (6) and blocked the tyrosine phosphorylation of a subset of host proteins (Fig. 2C) (6). We next determined if TepP is important for efficient bacterial replication by comparing the generation of infectious particles in cell lines infected with either wild type or Δ*tepP* mutants. A2EN cells, a newly derived cervical epithelial cell line (25), were particularly restrictive for the replication of Δ*tepP* stains as compared to HeLa cells (Fig. 2D). This defect appears to be unrelated to invasion as Δ*tepP* mutants were not impaired for entry into cells (data not shown).

**Figure 2:**
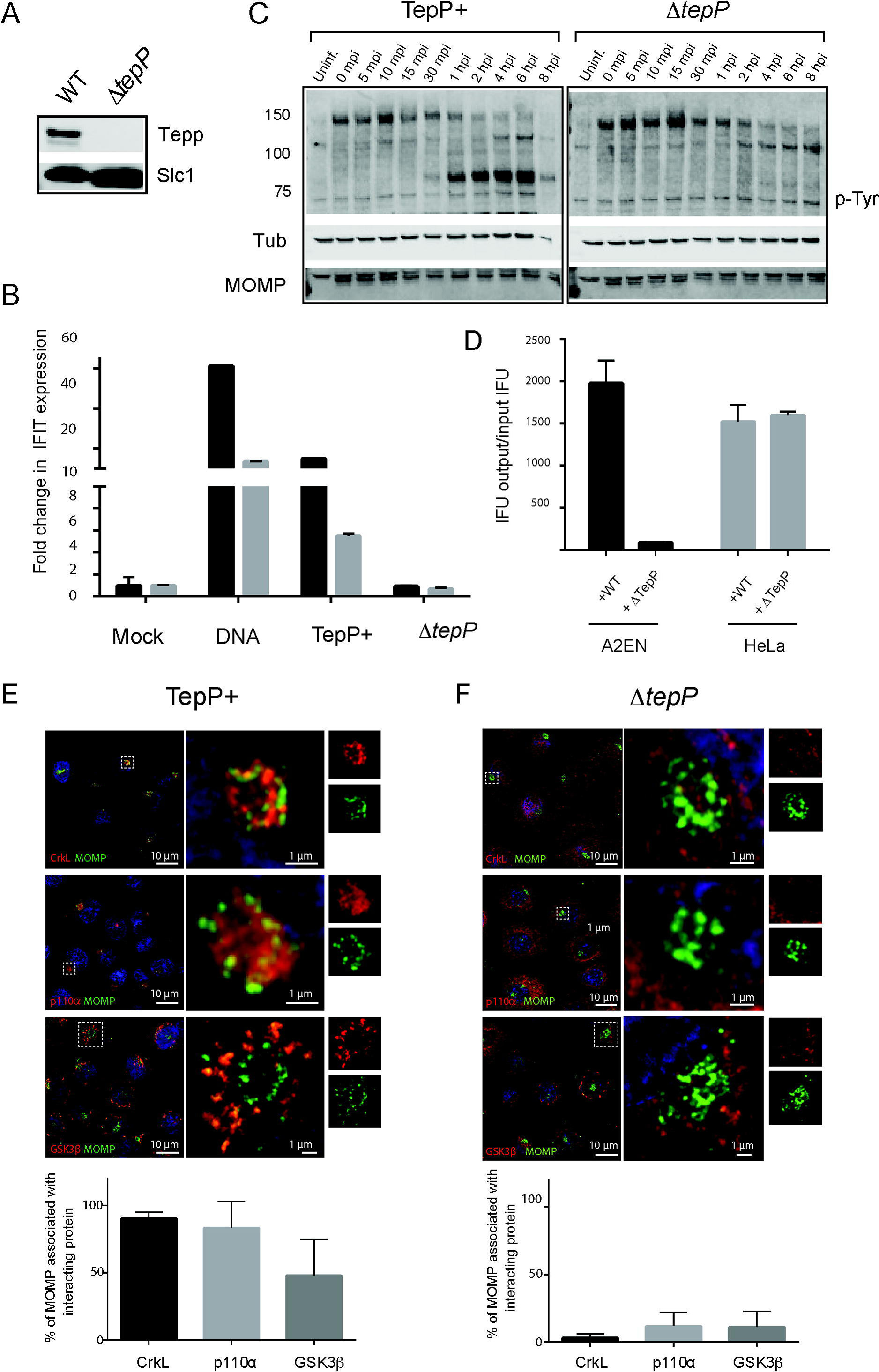
TepP is required for *C. trachomatis* replication in A2EN cells and to recruit CrkL and PI3K to nascent inclusions. **(A-D)** Phenotypic characterization of a *C. trachomatis* Δ*tepP::bla* insertional mutant. (**A**) Immunoblot analysis of TepP expression in HeLa cells infected for 24h with the indicated strains. Slc1: loading control. (**B**) A2EN cells were infected with CTL2 or Δ*tepP::bla* bacteria and the levels of *IFIT1* (black) and *IFIT2* (grey) transcripts (A) were assessed by quantitative RT-PCR. DNA transfection was used as a positive control for the induction of *IFIT* genes. **C**. Time course of appearance of major tyrosine phosphorylated (p-Y) proteins in A2EN cell infected with CTL2 or Δ*tepP::bla* insertional mutants. MOMP: bacterial outer major protein, Tub: tubulin. **D**. The replication potential of Δ*tepP::bla* mutants was assessed in HeLa and A2EN cells by the generation of inclusion-forming-units (IFU). **(E-F)** Subcellular localization of CrkL, PI3K (p110α) and GSK3β in *Chlamydia* infected cells. A2EN cells were infected with CTL2 or Δ*tepP::bla* bacteria for 4 hours and immunostained by anti-MOMP (green), anti-CrkL, anti-p110α, or anti-GSK3β (red) (**D**). Hoechst (blue) was used to detect DNA. Quantification of *Chlamydia* (MOMP) and p110α, CrkL, and GSK3β co-localization was performed on a single cell basis (n=20 and 30 cells/replicate). Student’s t-test identified there is a statistical difference between CTL2 and Δ*tepP::bla* bacteria with a p-value < 0.001.

We next tested if PI3K and CrkL are recruited to early inclusions where TepP translocation is most apparent (6). We infected A2EN cells with wild type or Δ*tepP C. trachomatis* and determined the extent of co-localization of TepP binding partners with intracellular bacteria by indirect immunofluorescence. Both PI3K (83%) and CrkL (90%) were strongly associated with clusters of intracellular bacteria that had migrated to a perinuclear region of host cells by 4 hpi (Fig. 2E). The accumulation of GSK at early inclusions was less prominent (48%). In contrast, we observed very little association of PI3K, CrkL or GSK with early inclusions formed by TepP-deficient *Chlamydia* (Fig. 2F). Because PI3K, Crk adaptors and GSK are involved in multiple signaling pathways and can impact the organization of the cytoskeleton, activation of innate immune factors, and expression of pro-survival signals (19, 21, 26, 27), our findings suggest that TepP recruits signaling proteins to influence signaling events.

### Src family kinases mediate the tyrosine phosphorylation of TepP but Src activity is not required for the recruitment of PI3K and CrkL to early inclusions

Because Tarp is phosphorylated by Src and Abl kinases (7, 8), we next tested if these kinases also play a role in the phosphorylation of TepP (7, 8, 28) by performing *in vitro* phosphorylation assays. Hexahistidine-tagged TepP purified from *E. coli* was incubated with ATP and cytosolic extracts derived from the following: Vero cells, Abl^-^/Arg^-^ mouse embryonic fibroblasts (MEFs), Src/Yes/ Fyn-deficient (SYF) MEFs, and their rescued counterparts. TepP was re-isolated and the extent of tyrosine phosphorylation assessed by immunoblot analysis with anti-phosphotyrosine antibodies. Recombinant TepP was phosphorylated after incubation with all cell lysates except for those derived from SYF MEFs, suggesting that Src-family kinases are required for TepP tyrosine phosphorylation *in vitro* (Fig. 3A). To determine if Src performs a similar function *in vivo* the same cells lines were infected with CTL2-M062G1 expressing TepP-FLAG. After IP with anti-FLAG antibodies, we assessed the extent of TepP tyrosine phosphorylation by immunoblot analysis (Fig. 3B). We observed diminished phosphorylation of TepP during infection of SYF MEFs, and increased phosphorylation upon overexpression of c-Src and to a lesser extent of Abl and Arg. Taken together, these findings suggests that the Src family kinases phosphorylate TepP during infection, although other kinases may also play an auxiliary role.

**Figure 3:**
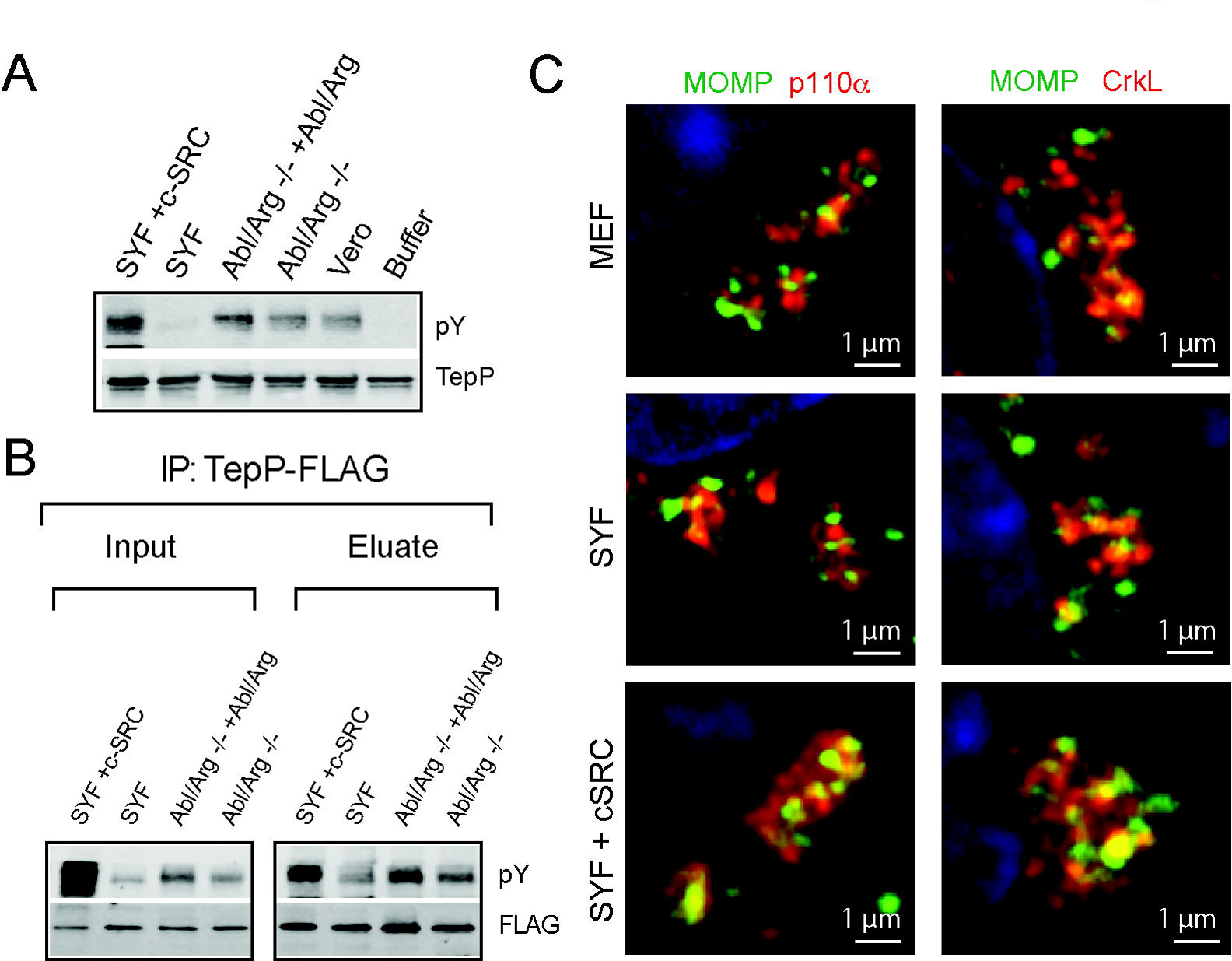
TepP is phosphorylated by Src kinases. **(A)** *In vitro* phosphorylation reactions were performed by incubating recombinant 6xHis-TepP with ATP and cell lysates derived from the indicated cell lines. The degree of TepP phosphorylation was assessed by immunoblotting with anti-pY antibodies after re-isolation of TepP on nickel beads. TepP is phosphorylated under all conditions except after incubation with SYF cell lysates and the buffer only control. **(B)** TepP-FLAG immunoprecipitations from cell lines infected with CTLM062G1 expressing TepP-FLAG. TepP phosphorylation was assessed after IP with anti-FLAG antibodies and immunoblotting with anti-pY antibodies. Note decreased levels of phosphorylation in Src, Yes and Fyn-deficient cells. **(C)** Mouse embryo fibroblast (MEF), SYF, and SYF+c-Src cell lines were infected for 4 h with *C. trachomatis* at an MOI of 20, fixed and stained. Infected cells were immunostained by anti-MOMP (green), anti-CrkL or anti-p110α (red), antibodies.

We hypothesized that Src-mediated phosphorylation of TepP would regulate the recruitment of SH2 and PTB domain proteins like PI3K (p85 subunit) and CrkL. We infected SYF and SYF+Src MEFs with wild type *C. trachomatis* and assessed the recruitment of PI3K and CrkL to nascent inclusions by immunofluorescence microscopy. Surprisingly, Src kinases were not required for the recruitment of PI3K or CrkL to inclusions (Fig. 3C). As a complementary approach, we generated phenylalanine substitutions at the three tyrosine (Y43 Y496 and Y504) identified as TepP phosphorylation sites (6). The TepP^Y43F/Y496F/Y504F^ variant was still capable of pulling down PI3K and CrkL and the mutant strain recruited these factors to early inclusions. However, this TepP variant was still tyrosine phosphorylated during infection (Fig. S1), suggesting that additional tyrosine phosphorylated residues in TepP may mediate interactions with host proteins.

Taken all together, this data suggests that while Src kinases are important for the full phosphorylation of TepP, these modifications do not appear to be essential determinant for TepP to interact with PI3K or CrkL in infected cells.

### PI3K association with TepP at early inclusions is independent of CrkL

PI3K and Crk adaptors are important signal transducers in growth factor mediated activation of receptor tyrosine kinases (29, 30). Both p85 and CrkL have SH3 and SH2 domains that mediate protein-protein interactions. Proteins that bind to Crk proteins at both the SH2 and SH3 domains include p85 itself, paxillin, C3G, Abl, Arg, Cas, and Sos (19, 22, 30-32). Similarly the p110 and p85 PI3K subunits interact with a variety of kinases including Fyn, Lyn, Src, Fak, and Bcr (33, 34). Because CrkL and PI3K can interact with each other (19, 22, 31) we tested if the association of these proteins with TepP is co-operative. We generated HeLa cell lines lacking the p110α subunit of PI3K, CrkL, or CrkI/II by CRISPR/Cas9-mediated gene disruption (Fig. 4A). These cells were infected with wild type *C. trachomatis* and the degree of co-localization of PI3K and Crk proteins with nascent inclusions was determined by indirect immunofluorescence. We did not detect any major differences in the efficiency of co-localization of the bacteria with PI3K in CrkL-deficient HeLa cells and vice versa (Fig. 4B), suggesting that CrkL and PI3K may engage TepP independently of each other. Similarly, CrkI/II-deficient cell lines were not impaired for the recruitment of PI3K to early inclusions (Fig. 4B).

**Figure 4:**
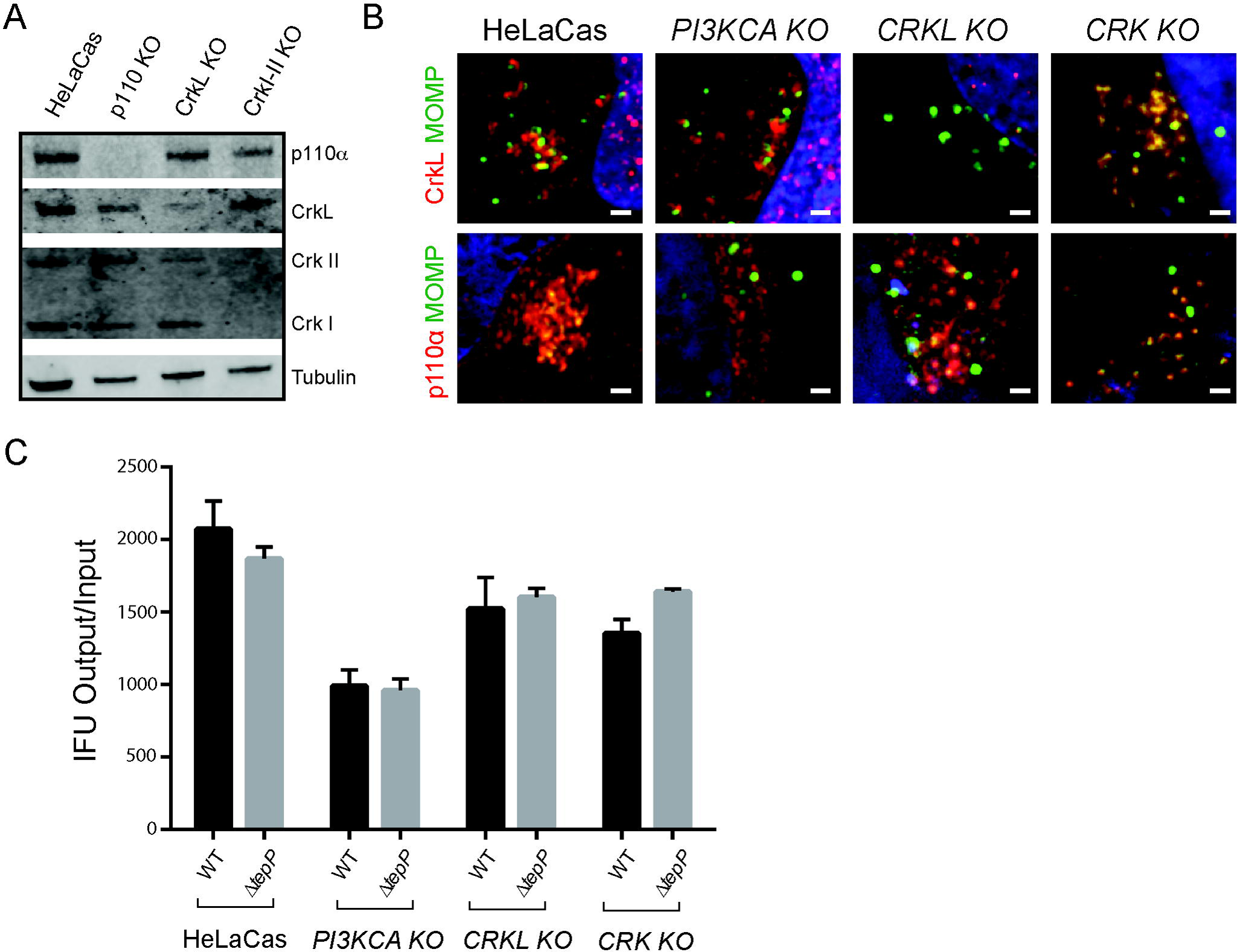
PI3K and CrkL independently associate with early inclusions. **(A)** HeLa cells stably expressing Cas9 (HeLa-Cas) were transduced with gRNAs specific to *PIK3CA*, *CRKL*, and *CRKI-II* to generated gene edited cell lines lacking expression of the respective target proteins. Loss of protein expression for each cell line was verified by immunoblot analysis using anti-p110α, anti-CrkL, and anti-Crk I/II. Tubulin levels were used as positive controls. There is a mild level of cross reactivity between CrkL and CrkII antibodies. **(B)** CrkL and p110α are recruited to early inclusions formed in *PI3KCA* and *CRKL* gene edited knock out cells (KO), respectively. HeLa-Cas cells and their edited derivatives were infected with CTL2 for 4h, fixed and immunostained with anti-p110α or anti-CrkL (red) antibodies. Host and bacterial DNA were detected with Hoechst (blue). **(C)** Replication of WT and TepP-deficient *C. trachomatis* in *PI3KCA*, *CRKL* and *CRK* gene edited knock out cells (KO)

Previous studies based on RNAi-mediated gene silencing suggested that both PI3K and Crk proteins are important for the replication of *Chlamydia* (35, 36). We assessed the efficiency of the various HeLa edited cell lines to generate *C. trachomatis* CTL2 infectious particles. PI3K-deficient HeLa cells displayed ∼50% defect in the generation of infectious EBs at 48 hpi. CrkL and CrkI/II deficient lines only displayed mild defects (<25%) (Fig. 4C). However, we did not observe differences in the generation of EBs between wild type and TepP-deficient *Chlamydia* among the various knockout HeLa cell lines, suggesting that the role of PI3K and Crk proteins in promoting *Chlamydia* replication, at least in HeLa cells, occurs independently of their interactions with TepP.

### PI3K modulates TepP-dependent Type I IFN responses

PI3K and CrkL participate in the activation of immune signaling (32, 37, 38). CrkL can interact with STAT5 to regulate the translocation of this transcription factor to the nucleus and the expression of Type I interferon-induced genes (32, 39). We tested if PI3K and Crk adaptors were required for the TepP-dependent induction of *IFIT* genes. PI3K, CrkL and Crk I/II deficient HeLa cells were infected with either wild type or Δ*tepP C. trachomatis* and *IFIT1* and *IFIT2* expression was assessed at 16 hours post infection by quantitative PCR. CrkI/II and CrkL-deficient HeLa cells did not display any significant differences in their response to either wild type or mutant *Chlamydia*. In contrast, PI3K deficient cell lines displayed an exaggerated *IFIT* response to wild type bacteria (Fig. 5). These cell lines also displayed a stronger response to cytoplasmic DNA, suggesting a broader role for PI3K activity in modulating response to cytosolic nucleic acids. Similar observations were made in HeLa cells treated with the PI3K inhibitor LY294002 in response to both DNA and *Chlamydia* infection (Fig. S2). Overall, these observations imply that one of the functions of p110α at nascent inclusions is to dampen Type I IFN response in response to TepP-expressing *Chlamydia*.

**Figure 5:**
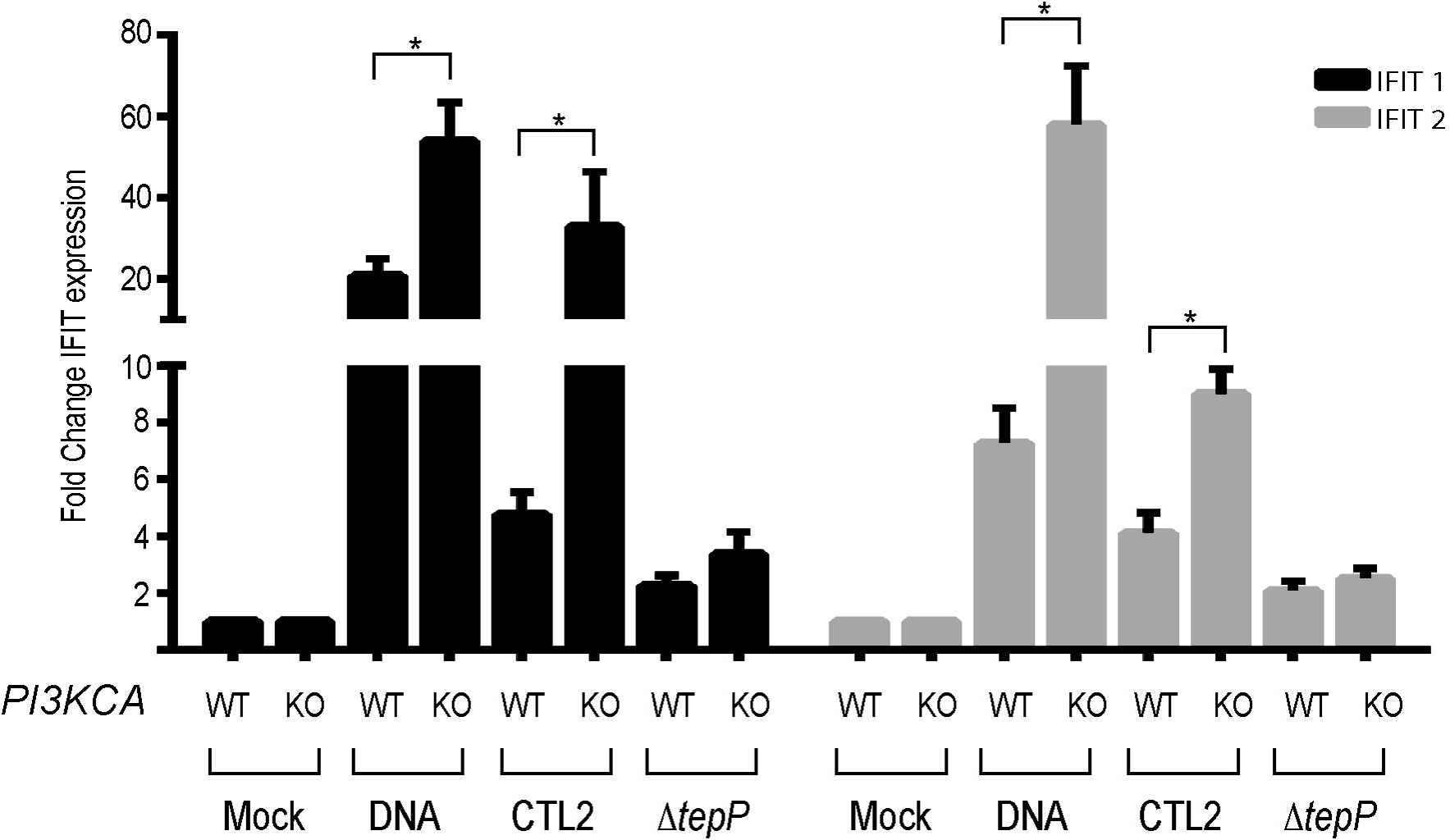
PI3K modulates TepP-dependent Type I IFN responses. HeLa-Cas and *PI3KCA* KO cells were infected with CTL2 or *ΔtepP* mutants and *IFIT1* (black bars) or *IFIT2* (grey bars) expression was assessed at 16 hpi by quantitative RT-PCR. WT and DNA samples for both *IFIT1 and IFIT2* are significantly different between HeLa-Cas and *PI3KCA* KO (p-value < 0.001, log-transformed anova with a priori contrasts). Fold chansge is calculated compared to mock infected isolates. DNA transfections were included as control for the induction of IFIT genes. The difference between HeLa-Cas and *PI3KCA* KO cells responses to DNA transfection or CTL2 infection is statistically significant (p-value < 0.01 Student’s t-test).

### TepP increases PI3K activity on early inclusions

In canonical PI3K/Akt signaling, activation of receptor tyrosine kinase (RTK) at the plasma membrane leads to the recruitment of PI3K and phosphorylation of phosphoinositide (4,5)-bisphosphate (PIP2) to generate phosphoinositide (3,4,5)-trisphosphate (PIP3) (21). Akt is then recruited to PIP3 enriched sites at the plasma membrane via its PH domain, leading to Akt phosphorylation and activation (21, 40). Activated Akt is an important regulator of multiple pro-survival signals (21, 27, 41). Upon *C. trachomatis* infection of HeLa cells, phosphorylation of Akt at Ser473 occurs with a bimodal distribution with early (<1h) and late (>12h) peaks of activation (42). We next tested if TepP contributed to the activation of PI3K and Akt. HeLa cells were infected with either wild type or Δ*tepP C. trachomatis* and the levels of Akt activation assessed by immunoblot analysis with anti phospho-Akt antibodies. We observed low levels of p-Akt levels at 4h post-infection and this activity was dependent on PI3K activity as the signal was blocked by pre-incubation with the inhibitor LY294002 (Fig. 6A). Infection with Δ*tepP* bacteria did not significantly alter in pAkt levels as compared to wild type *C. trachomatis* suggesting that PI3K activation at the plasma membrane is not overtly affected by TepP. The activation of Akt during *Chlamydia* infection is mostly mediated by the p110α isoform as no phospho-Akt signal was detectable in p110α KO cells (data not shown).

**Figure 6:**
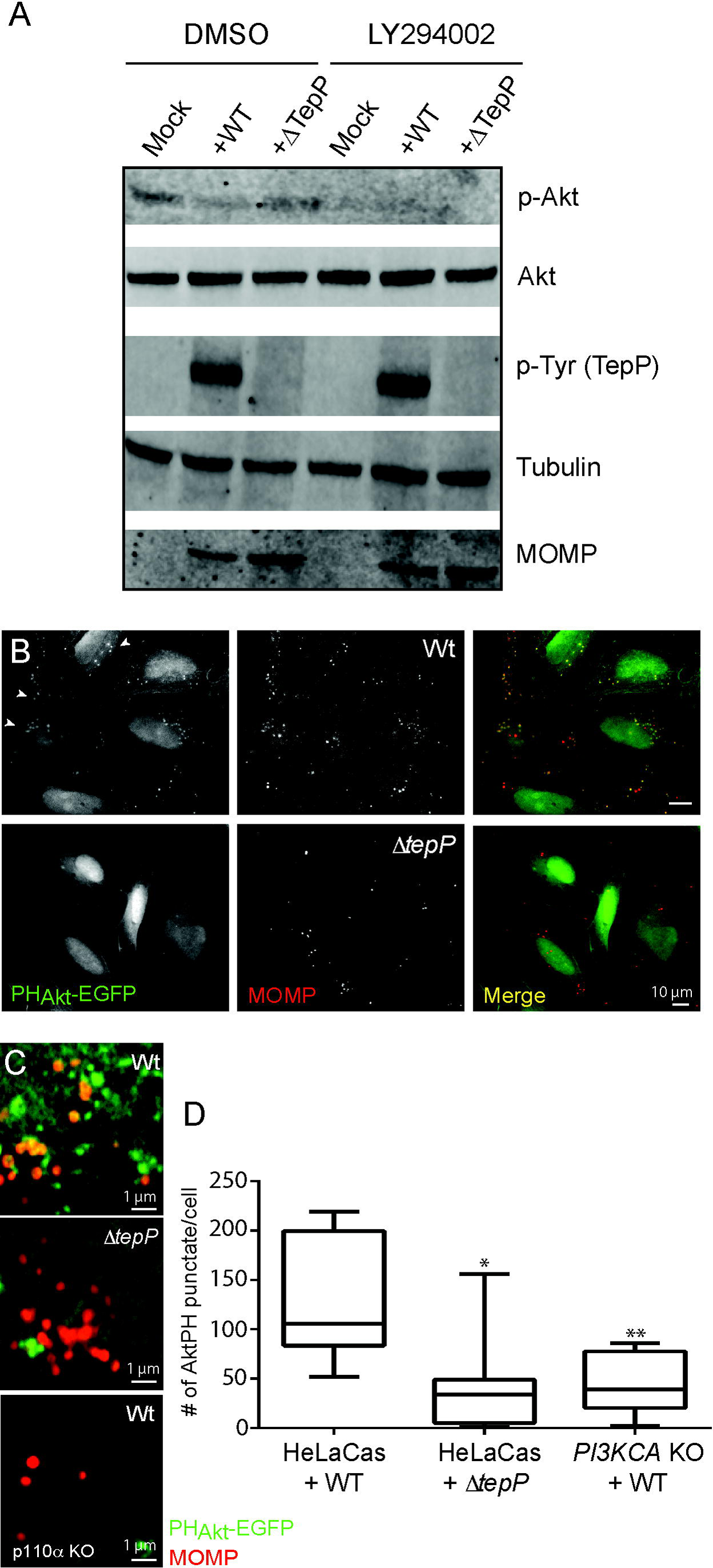
TepP activates PI3K at early inclusions. **(A)** Immunoblot analysis of HeLa-Cas cells infected with CTL2 or *ΔtepP::bla C. trachomatis* for 4h indicates that phospo-Akt levels are not significantly altered by TepP. **(B-D)** Accumulation of PIP3 positive puncta, as assessed by the recruitment of PH_Akt_-EGFP), in HeLa cells infected with CTL2 but not *ΔtepP::bla C. trachomatis*. HeLa cells were transfected with PH_Akt_-EGFP for 24h, and infected with the indicated *C. trachomatis* strains for 4h, fixed and immunostained with anti-MOMP antibodies (**B**). (**C**) Close up image of clusters of internalized *C. trachomatis* displaying association with PIP3 (PH_Akt_-EGFP positive). Bacteria associated PH_Akt_-EGFP intracellular puncta were significantly reduced in CTL2 infected *PI3KCA* KO HeLa cells. (**D**) Quantification of PIP3-positive puncta in cells infected with CTL2 or *ΔtepP::bla C. trachomatis* was performed on a per image basis with 7-10 fields total. Note that puncta formation required the p110α subunit of PI3K. Significance was determined by a Student’s t-test with * p< 0.01 and ** p< 0.001.

Although Akt phosphorylation is a common readout of increased activation of Class I PI3Ks, additional co-factors like PDK1 are required to phosphorylate Akt (21, 43). The most direct measurement of PI3K activity is the generation of PIP3. To assess the levels and spatial distribution of PIP3 pools in infected cells, we transfected cells with constructs expressing EGFP fusions to the PH domains of Btk and Akt that specifically bind to PIP3 (44). We observed an accumulation of PIP3 positive intracellular puncta in cells infected with wild type but not Δ*tepP C. trachomatis* (Fig. 6C-E). These puncta were observed in the vicinity of nascent inclusions but not always completely overlapping (Fig. 6D) and were not observed in PI3K deficient cells, suggesting that TepP mediates the localized formation of PIP3 though the activation of PI3K activity (Fig. 6E).

## Discussion

The process of *Chlamydia* entry into cells and establishment of a nascent inclusion is mediated by the controlled reorganization of the cytoskeleton and membranes by effector proteins. We determined that the early secreted effector protein TepP recruits the adaptor protein CrkL and the lipid kinase PI3K, leading to the localized synthesis of PIP3 on early inclusions (Fig. 1 and 2). Based on TepP’s ability to stably recruit PI3K, CrkL and CrkI/II proteins, and the role that Crk proteins play as adaptors linking receptors to signaling outputs, we hypothesize that TepP acts as a scaffold for the localized reprogramming of signaling proteins early during infection.

We generated a strain with *a bla* gene insertion in *tepP* in a clean genetic background to assess the phenotypic difference between wild type *C. trachomatis* and strains deficient only for the expression of TepP. All TepP dependent phenotypes described previously in a chemically derived *tepP* mutant (6) were validated using this new deletion strain and further allowed us to reveal a contribution of TepP to bacterial replication in A2EN cells. A2EN cells were recently derived from cervical tissue explants and maintain phenotypic and functional characteristics of endocervix epithelial cells, including cell polarization, mucus secretion, and expression of anti-microbial peptides in response to microbial TLR agonists (45).

Previous genome wide RNAi screens aimed at identifying host factors important for *Chlamydia* proliferation identified PI3K and Crk proteins as important for *Chlamydia* entry and replication (35, 46). These growth defects were confirmed in HeLa cells where p110α and Crk genes had been inactivated through CRISPR/Cas9-mediated gene editing, with p110α-deficient lines showing the greatest impairment (Fig 4). These host proteins may contribute to pathogen replication by facilitating entry, activation of pathways important for nutrient acquisition, and/or inactivation of cell defense mechanisms. However, although TepP is required for the recruitment of CrkL and PI3K to nascent inclusions early in infection, their contribution to enhancing bacterial replication in HeLa cells does not appear to occur through its interaction with TepP as Δ*tepP* mutants replicated to similar levels as wild type *C. trachomatis* in HeLa cells where Crk and PI3K had been inactivated (Fig. 4).

We expected that most host factors would specifically associate with phosphorylated versions of TepP. This appears not to be the case as the interaction between TepP and PI3K and CrkL, at least within the limits of our assays, was not affected by Src-mediated phosphorylation of TepP or by mutation of the three major tyrosines that are phosphorylated (Fig. 3 and S1). It is possible that other tyrosine kinases or additional tyrosine phosphorylation sites may independently contribute to the recruitment of PI3K, Crk adaptors, and/or other SH2 and PTB domain-containing proteins. Because Crk proteins bridge phosphorylated RTKs to p85 to activate PI3K (33), we expected that the binding of the p85/p110 complex to TepP would be mediated by CrkL. However, we find that recruitment of these proteins to early inclusions occurred independently of each other. The influenza viral protein NS1 independently binds to p85 and CrkL to activate PI3K activity (47, 48). TepP could be performing a similar function by assembling a tripartite Crk-TepP-PI3K complex that potentiates kinase activity as exemplified by the marked accumulation of PIP3 positive vesicles at the periphery of early inclusions. Alternatively, because the presence of TepP increased the efficiency of PI3K co-IP with CrkL (Fig 1C) there’s the possibility that Crk proteins further escalate the recruitment of additional p85/p110 to early inclusions and to enhance PI3K activity. We note that the activation of PI3K activity by TepP at intracellular sites did not lead to increased phosphorylation of Akt, suggesting that the activity of this kinase is uncoupled from events that may occur at the plasma membrane.

The prolonged recruitment of PI3K and Crk proteins on early inclusions that have migrated to perinuclear regions of infected cells correlates with the kinetics of TepP translocation after EB entry into cells (6). We predict that this recruitment and activation is independent of events that occur during the initial stages of infection (<1h) where engagement of RTKs like the EGFR (49), EphrRA2 (50) and PDGFR (35), and/or tyrosine phosphorylated Tarp may transiently recruit Crk proteins and PI3K to entry sites. The consequence of PI3K activation by TepP is the localized production of PIP3 at and in the vicinity of internalized bacteria (Fig. 6), which we predict lead to the recruitment of specialized PH domain-containing proteins that will further influence downstream signaling and/or cytoskeletal and membrane remodeling events. It remains to be determined how TepP orchestrates PI3K activity, Crk and GSK3 binding with the potential recruitment of additional factors. But the ultimate consequences of a failure to recruit all (or a subset of these factors) is a decrease in *Chlamydia* fitness in cervical epithelial cells and a decrease in the transcription of genes associated with Type I IFN responses.

Although PI3K and Crk proteins are important for full bacterial replication, as assessed by reduced growth in cells where these factors have been deleted, their role in promoting *Chlamydia* replication does not appear to be mediated by TepP, at least in HeLa cells. PI3K, but not Crk proteins, however, does appear to influence the magnitude of IFN responses to TepP-deficient bacteria (Fig. 5). PI3K knockout HeLa cells had a >5 fold in IFIT expression upon infection with TepP-deficient bacteria. PI3K regulates innate immunity in a variety of ways (48, 51) and inhibition of PI3K in dendritic cells enhances IFN-β transcription (52). It is possible that the absence of TepP increases the availability of microbial products from early inclusions to innate immune sensors or that PI3K activity modulates the function of sensors, which is consistent with the finding that basal expression of *IFIT* genes increases 2-3 fold in response to cytosolic DNA (Fig 5).

Overall, our findings begin to delineate important molecular components that are specifically assembled on *C. trachomatis* early inclusions that are ultimately important for fitness of the pathogen and for dampening of innate immune signaling in response to infection. The molecular players downstream of increased PI3K at intracellular sites and how they modulate the expression of immunity genes and bacterial survival in cervical epithelial cell remains to be determined.

## Materials and Methods

### Cell lines, bacterial strains and reagents

*Chlamydia trachomatis* serotype LGV-L2, strain 434/Bu (CTL2), and subsequently derived strains (Δ*tepP::bla*, CTL2-M062G1 (*tepP*^*Q103\**^), CTL2-M062G1 (pVec) (6), and CTL2-M062G1 (pTepP) (6)) were propagated in Vero cells maintained in Dulbecco’s Modified Eagle Medium (Sigma-Aldrich, St. Louis, Missouri, USA) supplemented with 10% fetal bovine serum (Mediatech, Manassas, Virginia, USA). Vero, HeLa (and associated Cas9 variants), and HEK293T cells were propagated in Dulbecco’s Modified Eagle Medium (Sigma-Aldrich, St. Louis, Missouri, USA) supplemented with 10% fetal bovine serum (Mediatech, Manassas, Virginia, USA). A2EN cells were propagated in keratinocyte-SFM medium (Gibco, Life Technologies corp., Grand Island, NY, USA) supplemented with 10 % FBS, 0.5 ng/mL human recombinant epidermal growth factor and 50 μg/mL bovine pituitary extract. EBs were purified by density gradient centrifugation using Omnipaque 350 (GE Healthcare, Princeton, New Jersey, USA) as previously described (18). PH-Akt-GFP was a gift from Tamas Balla (Addgene plasmid # 51465 (53)). Recombinant TepP was expressed in *Escherichia coli* strain BL21(DE3) as previously described (6). All reagents used are of analytical grade. The transfection reagent lipofectamine 2000 was purchased from Invitrogen. Opti-MEM serum-free media and rat tail collagen I were purchased from Gibco.

### Generation of *ΔtepP::bla C. trachomatis* strains

Recombinant *C. trachomatis* strains were generated through a modified version of the CaCl_2_ DNA transformation protocol previously described (54). TargeTron targeted gene disruption system (Sigma Aldrich,) was used to insert a *bla* cassette at the *tepP* locus, between amino acid 821 and 822 (ATCTCTCTGGATAATACAACGTCTGAGAAA - intron – TTGCTCATGTCCAGC) in CTL2. Plaque-purified recombinants were expanded in Vero cells and proper targeting was by confirmed by PCR with primers flanking the insertion site, whole genome sequencing, and by western blot analysis with anti-TepP antibodies (Fig. 2A).

### IFU burst assay

A2EN and HeLaCas cells were seeded onto 96 well plates (15,000 cells/well) and 24 h later, cells in each well were infected with the indicated strains (three biological replicates per time-point) at multiplicity of infection (MOI) of approximately 0.5. The number of input inclusion-forming units (IFU) was assesed at 24 hours by fixing infected cells with 100% Methanol (EMD Millipore) for 10 minutes on ice and immunostaining with anti-CTL2 sera and Alexafluor-conjugated secondary antibodies (Invitrogen Life Technologies, Carlsbad, California, USA). Images were acquired in a Zeiss Axioskop 2 upright epifluorescence microscope on at least 3 different fields per replicate and the number of inclusions were counted. To assess bacterial replication and infectivity, infections were allowed to proceed for 48h, cell were lysed by hypotonic lysis and IFUs calculated as described above on monolayers of Vero cells seeded onto 96 well plates. The total number of infected progeny released (output) was divided by the total number of input infectious particles. All statistical analysis was performed using Graphpad Prism.

### In vitro kinase Assays

His-TepP was purified from *E.coli* BL21 strain, grown in LB broth, with plasmid induction with IPTG (Sigma, 367-93-1) at a concentration of 0.5 mM for 4 hours at 37°C. Cells were lysed with lysis buffer (5M NaCl, 10% Triton-X100, 1M Imidazole, 250 mM PMSF, and 20 mM Phosphate) supplemented with Lysozyme (ThermoFisher Scientific, 89833), with sonication to reduce viscosity and centrifugation to pellet cell debris. Purification of His-TepP was preformed using nickel resin (ThermoFisher Scientific, 88221) and end-over-end incubation at 4°C for 1 hour. Resin was washed 3X with lysis buffer and subsequently washed 3X with *in vitro* kinase buffer (1M Hepes, 2M MgCl_2_, 250mM KCl, 1ATP, PMSF, Halt protease inhibitor cocktail (ThermoFisher Scientific, 78430), and cOmplete phosphatase inhibitor cocktail (Sigma, 4693159001)). Bound His-TepP was then incubated with whole cell lystates, lysed with *in vitro* kinase buffer and light sonication, from Vero, MEF, SYF, c+Src, Abl/Arg, Abl/Arg +Abl/Arg cells for 1 hour with end-over-end mixing at 4°C. Bound TepP was then washed 5X with *in vitro* kinase buffer without ATP. His-TepP was eluted from the nickel resin by incubation with Laemmli sample buffer and boiling. Levels of phosphorylation were assessed by immunoblot staining with anti-phosphotyrosine (Cell Signaling, 9411) and TepP levels assessed by immunoblot staining with anti-TepP (6).

### Indirect immunofluorescence microscopy

A2EN cells were seeded at a density of approximately 8 × 10^4^ cells/well on glass coverslips pre-coated with 30 μg/mL collagen in 20 mM acetic acid (Spectrum) for 5 min and rinsed twice with fresh media. HeLa and MEF cells were seeded at a density of 5 × 10^4^ cells/well on glass coverslips. The following day, CTL2 EBs were added at the indicated MOIs (20 or 100). Infections in A2EN cells were synchronized by centrifugation (500 × g for 5 min) at 10°C whereas infections in HeLa cells and MEFs were synchronized by centrifugation (1,000 × g for 20 min) at 10°C.The media was replaced and the infected cells transferred to a 37°C, 5 % CO_2_ humidified incubator. Coverslips were fixed with ice-cold 100% methanol for 15 min, rehydrated with PBS (3 × 5 min washes), and blocked with 2% BSA in PBS for 20 min

To assess the levels of PIP3 on early inclusions, HeLa cells were transfected with a plasmid encoding PH-Akt-GFP. After 24h, cells were infected with CTL2 or CTL2 *ΔtepP::bla* for 4h, rinsed twice with PBS and fixed with 1.5% formaldehyde (Sigma) in PBS for 20 min. Cells were quenched with 0.25% ammonium chloride (Sigma) and permeabilized/blocked with 2% BSA containing 0.1% saponin for 30 min.

Rabbit antibodies against TepP (1:50) (6), p110α (1:100) (Cell signaling #9411), Slc1 (1:500) (6), Crk (1:50) (BD Transduction Laboratories 610035 clone 22), CrkL (1:100) (Thermo Fisher Scientific PA5-28622) and a mouse antibody against MOMP (1:500)(sc-57678, Santa Cruz) were diluted in PBS containing 2% BSA. Secondary antibodies include Goat-anti-Mouse (H+L) Alexa Fluor 488 and Alexa Fluor 555 (Thermo Fisher Scientific A11001 and A21422), and Goat-anti-Rabbit (H+L) Alexa Fluor 488 and Alexa Fluor 555(Thermo Fisher Scientific A11034 and A21428), were diluted in PBS containing 2% BSA. Hoechst (Invitrogen; H3570) was diluted in PBS and incubated on the cells for 10 min. Coverslips were mounted on slides using Fluorsave mounting media (CalBiochem).

Cells were imaged using a confocal laser scanning microscope (LSM 880; Zeiss)d equipped an Airyscan detector (Hamamatsu) and diode (405 nm), Aragon ion (488 nm), double solid-state (561 nm), and helium-neon (633 nm) lasers. Images were acquired using a 60x/1.4 NA oil objective (Zeiss) and deconvolved using automatic Airyscan Processing in the Zen Software (Zeiss). In Fig 6B, PH-Akt-GFP expressing HeLa cells were imaged using the Zeiss AxioObserver Z.1 widefield microscope equipped with a motorized stage and a camera (Axiocam MRm; Zeiss). Image were acquired using a 63x/1.4 NA oil objective (Zeiss) and the AxioVision 4.1 software.

### Image Processing

All images were processed using ImageJ open source software (ImageJ). Linear adjustments were made for all images and max projections from Z-stacks are portrayed. Quantification of recruitment of proteins to the nascent inclusion was performed in ImageJ using the JACoP (Just another co-localization plugin) on a per cell basis on each z-slice of an image. The percent co-localization was calculated using distance based co-localization and the % of positive thresholded pixels of MOMP that associate with thresholded pixels of the either p110α, CrkL, or GSK3β proteins. Thresholds were set to eliminate background signals in both MOMP and TepP associated protein channels. Strictly set thresholds help to reduce the number of false positive co-localization calculations. Quantification of PIP3-positive puncta (PH-Akt-GFP) was performed by using the *Analyze Particles* function in ImageJ. Strict thresholding parameters were applied to reduce the number of false positives.

### Immunoprecipitations

Antibodies against CrkL (PA5-28622), p110α (Cell Signaling Technology 4249S (α)), and FLAG (conjugated to magnetic beads (Sigma-Aldrich, St. Louis, Missouri, USA M8823) were incubated for 2 hours at 4°C with total cell lysates derived from 6x10^6^ A2EN cells infected with the indicated *C. trachomatis* CTL2 strains at an MOI of 100. Infected cells were lysed in 750 μL of Radioimmunoprecipitation assay buffer (RIPA) buffer (Sigma R0278) supplemented with 1X EDTA free protease inhibitor cocktail (Roche, Basel, Switzerland) and Halt phosphatase inhibitor (Thermo Risher Scientific 78428). After a 2-hour period, CrkL and p110α samples were incubated for 1 hour at 4°C with magnetic protein A conjugated beads (SureBeads Protein A Magnetic Beads, Bio-Rad 161-4011). All magnetic beads slurries were then washed 5X with 200 μL of lysis buffer and bound proteins were eluted in Laemmli sample buffer and identified by western blot analysis.

### Identification of TepP-FLAG interacting proteins

Four 15 cm dishes of confluent human A2EN cells were infected either with CTL2-M062G1 (*tepP*^*Q103\**^) or CTL2-M062G1 transformed with a plasmid expressing TepP-FLAG. Cells were infected at an MOI of 50 by rocking at 4°C for 30 min in HBSS (Hank’s Balanced Salt Solution) (Invitrogen Life Technologies, Carlsbad, California, USA), following by shifting to 37°C after adding warm keratinocyte-SFM medium. After 4 hours, cells were washed once with cold PBS and lyzed on ice by scrapping in lysis buffer (25 mM Tris, 150 mM NaCl, 1 mM EDTA, 1 % NP-40, 5 % glycerol; pH 7.4) supplemented with 1 mM phenylmethylsulphonyl fluoride (PMSF), 1X EDTA-free protease inhibitor cocktail (Roche, Basel, Switzerland) and Halt phosphatase inhibitor (Pierce, Rockford, Illinois, USA). Cell debris was pelleted by centrifugation and the supernatants were transferred to a new tube. Around 100 μl of anti-FLAG M2 magnetic beads (M8823) were added to each tube and mixed by constant rotation at 4°C for 4 hours. After 5 washes with lysis buffer, the bound proteins were eluted with Pierce Elution buffer (pH 2.8) (Pierce, Rockford, Illinois, USA)

Proteins were concentrated using 0.5 mL Amicon 10 MWCO filters into ammonium bicarbonate. After concentration, samples were separated on a NuPAGE 4-12% Bis-TRIS SDS-PAGE gel (ThermoFisher Scientific). After staining with Novex colloidal coomassie stain (ThermoFisher Scientific), three bands were isolated for each sample, covering the approximate molecular weight range from 20 kDa to 150 kDa. The three bands per sample were excised, de-stained, and the proteins in the bands were digested with trypsin according to the ‘‘In-Gel Tryptic Digestion Protocol’’ available at (http://www.genome.duke.edu/cores/proteomics/sample-preparation/). Briefly, bands were destained with 1:1 MeCN/water, reduced with 10 mM dithiothreitol and alkylated with 20 mM iodoacetamide, then dehydrated in MeCN and swelled in 50 mM ammonium bicarbonate containing 10 ng/ml trypsin. Digestion was carried out overnight at 37°C, and digestion was quenched and peptides were extracted using 0.1% v/v TFA in 1:1 MeCN/water. Samples were dried and reconstituted in 10 ml 1/2/97 v/v/v TFA/MeCN/water for mass spectrometry analysis. Liquid chromatography-tandem mass spectrometry (LC-MS/MS) for peptide sequencing was performed on a nanoAcquity UPLC coupled to a Synapt G2 HDMS (Waters Corporation). Raw data was processed in Mascot Distiller (Matrix Sciences) and Mascot Server v2.5 (Matrix Sciences) was used for database searching. A custom *.fasta database was constructed from combining the curated human proteome (uniprot.org) and *Chlamydia trachomatis* L2/434/Bu genome along with common laboratory contaminants. Database searching used 10 ppm precursors and 0.04 Da product ion tolerance, fixed carbamidomethylation (Cys), variable oxidiation (Met) and deamidation (NQ). Database search results were curated in Scaffold (Proteome Software) to a 0.38% peptide FDR and 0.5% protein FDR, using decoy database searching and the PeptideProphet algorithm.

### RNA isolation and RT-qPCR

RNA was collected from 3 wells of a 6-well plate of infected cells at 16 hpi using the RNeasy plus mini kit (Qiagen, 74134). A2EN, HeLa-Cas9, and HeLa-Cas9 KO cell lines were seeded 24 hours prior to infection at a density of approximately 0.8 × 10^6^ cells per well. The cells were then infected with gradient purified EBs of either CTL2 or CTL2 Δ*tepP::bla* strains at an MOI of 10. As controls, cells were also mock infected or transfected with 5ug of a linearized bacterial plasmid (pET-24d), using jetPRIME transfection system (Polyplus-transfection, VWR 89129-922). RT-qPCR was conducted using the Power SYBR Green RNA-to-CT™ 1-Step Kit (Thermo-Fisher Scientific). To assess differential gene expression of the *IFIT1-2* (primers described in (6)) relative expression levels were determined according to the comparative CT method (55) using β-actin mRNA as reference for normalization. Mock, DNA, WT, and Δ*tepP* treatments were analyzed via log-transformed anova (aov function, “stats” package) in R (56) with a priori contrasts between HeLa-Cas9 and *PI3KCA* KO cell lines. WT and DNA both showed significant (p<0.001) differences between HeLa-Cas9 and *PI3KCA* KO cell lines while there were no significant differences for Mock or Δ*tepP* treatments between HeLa-Cas9 and *PI3KCA* KO. This was observed for both *IFIT1* and *IFIT2*.

### Generation of CRISPR/Cas9 KO cells

HeLa KO cell lines were generated by stable integration of vectors expressing single guide (sgRNA) into Cas9 expressing HeLa cells. The following sgRNAs were used: GAGGACATGGTGTTGGACCG (*CRKL*), GACTTTAGAATGCCTCCGTG (*PI3KCA*), GGGGAGGTTGAGTCGGCAGG (*CRK*). To generate transducing viruses, HEK293T cells were co-transfected with the vectors; pasPAX2, pCMV-VSV-G, and the sgRNA in lentiGuide-Puro (addgene), three times at 12 hour intervals. Supernatants containing virus were collected from, filter sterilized (0.2 μm), and incubated with HeLa-Cas9 cells (Duke Functional Genomics Core facility) at 48 and 72 hours. At 24 hours after the last round of viral infections, the media was replaced with fresh media containing 5 μg/mL of puromycin to select for stably transduced cells. Selection was continued until all cell in mock virally infected cells were dead. Cells were then clonally isolated by limiting dilution and ach cell line was assessed for protein expression by immunoblot analysis.

## Supplemental Figure Legends

**Table S1: Summary of proteins identified by LC-MS/MS as potential binding partners for TepP**

**Supplemental Figure 1: The mutant TepP^Y43F/Y496F/Y504F^ is tyrosine phosphorylated and capable of recruiting CrkL and p110**α **to early inclusions**

**(A)** A2EN cells infected with CTL2-M062G1(pVec), CTL2-M062G1(pTepP), and CTL2-M062G1(pTepP^Y43F/Y496F/Y504F^) for 1, 2, and 4 h. Immunoblot analysis with anti-pY and anti-FLAG indicate levels of recombinant protein expression and tyrosine phosphorylation.

**(B)** A2EN cells were infected with CTL2-M062G1 (pTepP^Y43F/Y496F/Y504F^) for 4h, fixed and immunostained for *Chlamydia* with anti-MOMP antibodies(green). CrkL and p110α (red)were detected with specific antibodies. Nucleic acids were stained with Hoechst (blue).

**Supplemental Figure 2: Treatment with the PI3K inhibitor LY294002 increase *IFIT* gene expression upon infection.**

*IFIT* expression is induced in HeLa-Cas upon infection with CTL2 for 16h or upon DNA transfection. Treatment with LY294002 increase IFIT expression upon DNA transfection or infection with CTL2, but not after infection with Δ*tepP::bla* mutants. Fold change is calculated compared to mock infected isolates. WT and DNA samples for both *IFIT1 and IFIT2* are significantly different between HeLa-Cas and *PI3KCA* KO (p-value < 0.001, log-transformed anova with a priori contrasts).

## Supplementary Materials and Methods

### Generation of TepP point mutants

TepP tyrosine phosphorylation mutants were generated using site directed mutagenesis with the Q5 site-directed mutagenesis kit (New England Biolabs, E0554S). Each tyrosine site (Y43, Y496, and Y504) was mutated individually to phenylalanines in the +pTepP vector (6). Each version of the plasmid was transformed into *C. trachomatis* L2-M062G1 (TepP deficient strain) using a modified version of the CaCl_2_ transformation protocol previously described (54). Expression and translocation of TepP variants was verified with FLAG anti-bodies in immunoblot and immunofluorescence staining.

### LY249002 Inhibitor treatment, RNA isolation, and RT-qPCR

HeLa-Cas cells were seeded 24 hours prior to infection at a density of approximately 0.8 × 10^6^ cells per well. Cells were incubated with DMSO (solvent only control) or the PI3K kinase inhibitor LY294002 (Cell Signaling, 9901S) at a concentration of 20μM/mL for 1 hour prior to infection. Cells were then infected with gradient purified CTL2 or CTL2Δ*tepP::bla* EBs at an MOI of 10. As a control, cells were also mock infected or transfected with 5ug of a linearized bacterial plasmid (pET-24d) using jetPRIME transfection system (Polyplus-transfection, VWR 89129-922). RNA was collected from three wells of a 6-well plate at 16 hpi, DMSO or inhibitor treatment was maintained throughout infection. RT-qPCR was conducted using the Power SYBR Green RNA-to-CT™ 1-Step Kit (Thermo-Fisher Scientific). To assess differential gene expression of the IFITs (primers described in (6)) relative expression levels were determined according to the comparative CT method, using β-actin mRNA as reference for normalization. Mock, DNA, WT, and ΔTepP infections were analyzed via log-transformed anova (aov function, “stats” package) in R (56) with a priori contrasts between DMSO and LY294002 treated HeLa-Cas cells. WT and DNA both showed significant (p<0.0001) differences between DMSO and LY294002 treated HeLa-Cas cells while there were no significant differences for Mock, and small difference (p<0.05) during ΔTepP infection between DMSO and LY294002 treated HeLa-Cas cells. This was observed for both *IFIT1* and *IFIT2*.

## Acknowledgements

We thank the Duke University Proteomics Core and the Duke Functional Genomics Core facilities for their technical support. This work was supported by NIH awards AI 100759 and AI123083.

